# mcPBWT: Space-efficient Multi-column PBWT Scanning Algorithm for Composite Haplotype Matching

**DOI:** 10.1101/2022.02.02.478879

**Authors:** Pramesh Shakya, Ardalan Naseri, Degui Zhi, Shaojie Zhang

## Abstract

Positional Burrows-Wheeler Transform (PBWT) is a data structure that supports efficient algorithms for finding matching segments in a panel of haplotypes. It is of interest to study the composite patterns of multiple matching segments or blocks arranged contiguously along a same haplotype as they can indicate recombination crossover events, gene-conversion tracts, or, sometimes, errors of phasing algorithms. However, current PBWT algorithms do not support search of such composite patterns efficiently. Here, we present our algorithm, mcPBWT (multi-column PBWT), that uses multiple synchronized runs of PBWT at different variant sites providing a “look-ahead” information of matches at those variant sites. Such “look-ahead” information allows us to analyze multiple contiguous matching pairs in a single pass. We present two specific cases of mcPBWT, namely *double-PBWT* and *triple-PBWT* which utilize two and three columns of PBWT respectively. *double-PBWT* finds two matching pairs’ combinations representative of crossover event or phasing error while *triple-PBWT* finds three matching pairs’ combinations representative of gene-conversion tract.

## 1 Introduction

Durbin’s PBWT (positional Burrows-Wheeler Transform) [1] is an efficient data structure that operates on a panel of haplotypes that are bi-allelic to find all the matching segments of a given minimum length threshold, *L*. In addition to that, it’s also capable of finding matches for a query haplotype against a reference haplotype panel and other useful compression algorithms.

Given the linear time complexity of PBWT algorithms, they scale well to large datasets. Because of this, they have widely been incorporated in state-of-the-art statistical phasing and imputation tools [2, 3, 4]. PBWT algorithms have also been used to find identical-by-descent (IBD) [5, 6, 7] segments shared among individuals of a population. IBD segments are segments of chromosome shared among individuals such that they share a most recent common ancestor. Numerous features of IBD segments including their counts, length distribution, etc have been studied as they reveal useful information of the population history, selection pressure, and the disease loci [8]. Many other variations of the PBWT algorithm have been developed that tackle variety of problems. gPBWT [9] provides a method to efficiently query graph-encoded haplotypes, d-pbwt [10] provides efficient algorithms for query haplotype insertion and deletion, mPBWT [11] provides algorithms to deal with multi-allelic panels, [12] allows wildcard characters in the PBWT panel to study relative fitness of genomic variants, and cPBWT [13] and related works [14, 15] extend pairwise segment matching to multi-way matching, i.e., clusters of haplotype matches.

It is of interest to study composite haplotype matching patterns. For example, two long segment matches of a haplotype that are adjacent to each other may indicate a recombination event or an error of the phasing method. Another example is a combination of three segment matches, two long ones between the same pair of haplotypes, surrounding a relatively short one in the middle, that is a hallmark of a gene conversion. However, most existing problem formulations of PBWT algorithms are to find single matching segments between pairs of haplotypes or a single matching block among a cluster of haplotypes.

Recently, bi-directional PBWT [16] (or bi-PBWT) was the first to study composite matching patterns of more than one matching blocks. bi-PBWT finds all matches between sufficiently long matching blocks at both sides of a site, with a small gap of tolerance. The bi-PBWT algorithm is a two-pass scanning algorithm, first scanning backwards, storing the reverse PBWT data structure, and then a second pass scanning forward and makes the block matching. We wish to generalize the two-pattern matching problem to more, and with a more general definition of the connections between individual matching patterns.

Here, we formulate the problem of composite haplotype matching. Conceptually, the goal of composite haplotype matching problem is to find a number of pairwise matching segments or matching clusters, each one is long enough, with small enough gaps/overlaps, and the haplotype IDs of these segments satisfy certain condition (e.g., having a single id shared with all segments, and the other IDs may or may not belong to the same individual). The phasing error pattern, recombination, and the gene conversion each can be seen as special cases of such composite haplotype matching patterns.

In this paper, we introduce a space-efficient algorithm, mcPBWT, that utilizes two or more synchronized scans over multiple columns of PBWT to compare and analyze multiple sets of matches. While a naive solution in the style of bi-PBWT that stores pre-computed PBWT is time-efficient, the space-efficiency of pre-computed panel can be inconveniently large for biobank-scale data. Our algorithm’s multi-column idea allows various columns of PBWT to exchange information and integrate multiple single matches without a large memory-footprint or disk-usage. The algorithm also makes a single pass of a haplotype panel. This online nature and generalizability of the algorithm will provide an efficient way of studying complex set of matches.

## 2 Preliminaries

### 2.1 PBWT Overview

PBWT (positional Burrows-Wheeler Transform) is an efficient data structure that finds all matches of user-specified minimum length (*L*) in an efficient manner given a panel of haplotype sequences. In his paper, Durbin [1] defines a *haplotype panel X* as a set of *M* haplotype sequences *x*_*i*_ ∈ *X, i* = 0, 1, 2, …, *M* − 1. Each sequence *x*_*i*_ has *N* SNP sites indexed by *k, k* ∈ {0, 1, …*N* − 1}. All the sites are assumed to be bi-allelic, namely *x*_*i*_[*k*] ∈ {0, 1}. *Locally maximal match* is defined as a match between two haplotype sequences *s* and *t* from *k*_1_ to *k*_2_ such that, *s*[*k*_1_ − 1] ≠ *t*[*k*_1_ − 1], or *k*_1_ = 0 and *s*[*k*_2_] ≠ *t*[*k*_2_], or *k*_2_ = *N*. A match is a *long match* if it is locally maximal and its length satisfies a user-specified length threshold. *Prefix array a* contains *N* + 1 reverse prefix sorted orderings of the sequences, one for each *k* ∈ 0…*N*. It can also be thought of as a permutation of indices of the haplotype sequences that range from 0 to *M* − 1 for every *k* ∈ {0, 1, 2…*N*}. *a*_*k*_ is the *k*-th reverse sorted ordering of the haplotype sequences up to the site *k* − 1. In any *a*_*k*_, adjacent sequences are maximally matching until *k*. 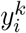 is the *i*-th sequence in *a*_*k*_, 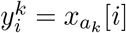. The divergence array *d*_*k*_ stores the starting position of locally maximal matches ending at *k* between a sequence and the preceding sequence in *a*_*k*_.

### 2.2 Composite Haplotype Matching

Here, we generalize the notion of haplotype matching in a panel. A *single haplotype match pattern* (or single match) in a haplotype panel is defined as *p* = (*c, k*_1_, *k*_2_), where *c* is a subset of the total set of haplotype indexes *C* = {0, …, *M* − 1}, and the haplotype sequences match between sites *k*_1_ and *k*_2_: *x*_*i*_[*k*_1_, *k*_2_) = *x*_*j*_[*k*_1_, *k*_2_), for any *i, j* ∈ *c*. Here, the *length* of the pattern is *L*(*p*) = |*k*_2_ − *k*_1_|, and the *width* of the pattern is *W* (*p*) = |*c*|. We can also denote the *sequence id set* of *p* as *c*(*p*) = *c*, the *left boundary* of *p* as *l*(*p*) = *k*_1_, and the *right boundary* as *r*(*p*) = *k*_2_. In general, *c* 2 indicates a cluster of haplotypes matching. For pairwise matching, |*c*| = 2. The problem of single pattern haplotype matching is, given a predefined length cutoff *L* and width cutoff *W*, find all patterns *p*, such that *L*(*p*) ≥ *L* and *W* (*p*) ≥ *W* in a haplotype panel.

Further, for two single haplotype match patterns *p* and *q*, we define *q g-follows p* if they are adjacent, i.e., the *gap* (or *overlap)* between them, *g*(*p, q*) = *l*(*q*) − *r*(*p*), is small: |*g*(*p, q*)| ≤ *g* and some haplotypes are shared among their sequence id sets *c*(*p*) ⋂ *c*(*q*) ≠ ∅.

With that, we define a *composite haplotype match pattern* in a haplotype panel as a series of *B* single matches, 𝒫 = {*p*_*b*_}, *b* = 0…*B* − 1, that *g*-follow each other, i.e., *p*_*i*_ *g*-follows *p*_*i−*1_, for *i* = 1..*B* − 1, and they share some common haplotypes 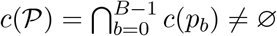. We call *B* the *span* of 𝒫, and *c*(𝒫) the *thread* of 𝒫.

The problem of composite haplotype matching pattern is, given a predefined set of length cutoffs {*L*_*b*_}, *b* = 0…*B* − 1, width cutoff *W*, gap tolerance *g*, the span *B*, the thread width *w*, find all composite patterns 𝒫 = {*p*_*b*_}, *b* = 0..*B* − 1 such that *L*(*p*_*b*_) ≥ *L*_*b*_, *W* (*p*_*b*_) ≥ *W, p*_*i*_ *g*-follows *p*_*i−*1_, for *i* = 1..*B* − 1, and |*c*(𝒫) | = *w*. Of course, it is possible to specify different width cutoffs and gap tolerances for individual single match. We omit that for simplicity of presentation.

In this work, we mainly focus on *double haplotype match patterns* (*B* = 2) or *triple haplotype match patterns* (*B* = 3). We will also mainly focus on pair segment matching (*W* = 2), and single thread composite patterns (*w* = 1). We will present memory-efficient multi-column scanning PBWT-based algorithms: *double-PBWT* for identifying double haplotype match patterns, and *triple-PBWT* for triple haplotype match patterns.

## 3 Multi-column PBWT

In PBWT, a single column scans the haplotype panel from left to right and updates divergence values and prefix arrays to output long matches. Let *P*_*k,L*_ signify the active single column of PBWT operating at site *k* finding matches of length at least *L* **ending** at site *k*. If the panel were to be scanned in a reverse fashion from right to left then, *R*_*N−k,L*_ would signify the active single column of PBWT operating at site *N* − *k* finding matches of length at least *L* ending at site *N* − *k*. This would allow the user to compare two sets of matches at site *k* (or *N* − *k* if looking from right to left) i.e. matches from *P*_*k,L*_ and *R*_*N−k,L*_. This method would require the panel to be scanned twice if we wanted to compare such matches along different SNP sites of the panel. In fact, one approach would be to scan the panel in reverse and save all the precomputed divergence values and prefix arrays before hand as in [16]. While this method works well, it occupies disk-space and needs to be loaded into memory which might not be efficient for larger panels. In contrast, mcPBWT is capable of providing the same information on a single pass of the haplotype panel from left to right using multiple columns of PBWT without having to pre-scan the panel and save the values on disk.

The general idea of mcPBWT is to utilize the information obtained from “look-ahead” PBWT columns. Here, a new PBWT column operating at site *k* + *L* that finds all the matches **starting** at site *k* is denoted by *P*′_*k*+*L,L*_ such that matches found by columns *P*′ and *R* are the same i.e. *P*′_*k* +*L,L*_ = *R*_*N−k,L*_. So mcPBWT would consist of a set of PBWT columns where the columns are finding matches ending at a site or starting few sites before. Even though, we only talk about two columns of PBWT here, similar approach can be also used for the case of three columns of PBWT. In fact, we can spawn multiple such columns of PBWT and at each column, the user has the flexibility to find either matches ending at that site as in PBWT or have the flexibility to find matches starting certain number of sites before, depending on the use case. First, we discuss the divergence value properties that enable us to find the matches **starting** at a certain site and then the two specific cases of mcPBWT in the following sections.

### 3.1 Divergence Value Properties

There are two major properties of the divergence values that assist in finding the matches starting at a certain site efficiently. The first property asserts that in a divergence array, adjacent divergence values cannot be equal unless it is zero (or when a match does not exist)[1], while the second property that extends the first property asserts that between any two consecutive equal divergence values there must be a divergence value that is greater than those equal values. These properties are presented formally as two lemmas below.

#### Lemma 1.

*Two adjacent divergence values aren’t equal unless it is zero (or when the divergence value greater than current k, i*.*e. when there is no match)*.

*Proof*. This property mentioned by Durbin, asserts that *d*_*k*_[*i* − 1] ≠ *d*_*k*_[*i*], 0 *< i < M* except when *d*_*k*_[*i*] = 0 or *d*_*k*_[*i*] = *k*. For any index *i*, the divergence value at some site *k, d*_*k*_[*i*] gives the starting position of a match between haplotypes at index *i* and *i* − 1. Since the panel is bi-allelic and reverse prefix sorted, this means that *y*_*i−*1_[*d*_*k*_[*i*] − 1] must be 0 and *y*_*i*_[*d*_*k*_[*i*] − 1] must be 1. The same condition holds for *d*_*k*_[*i* − 1] in that *y*_*i−*2_[*d*_*k*_[*i* − 1] − 1] must be 0 and *y*_*i−*1_[*d*_*k*_[*i* − 1] − 1] must be 1. But if we assume that *d*_*k*_[*i*] = *d*_*k*_[*i* − 1], there is a contradiction on the value for *y*_*i−*1_[*d*_*k*_[*i*] − 1]. This proves that the adjacent divergence values can’t being equal unless the divergence value is equal to 0 or *k* (match does not exist). □

#### Lemma 2.

*In a divergence array d*_*k*_, *assume d*_*k*_[*i*] *= x, and there exists another index g, g < i and g* ≠ *i* − 1 *where g is the first index preceding i to have divergence value equal to x i*.*e. d*_*k*_[*g*] *= x, then there must be a divergence value greater than x at index h, where g < h < i and d*_*k*_[*h*] *> x (except when x* = 0*) for* 0 *< g, h, i < M*.

*Proof*. This property is essentially an extension of lemma 1. In a bi-allelic panel, *d*_*k*_[*i*] = *x* asserts that *y*_*i−*1_[*x* − 1] = 0 and *y*_*i*_[*x* − 1] = 1. For index *h* in the range (*g, i*), the divergence value can either be *d*_*k*_[*h*] *< x* or *d*_*k*_[*h*] *> x* since *g* is the first index preceding *i* where *d*_*k*_[*g*] = *x*. If we assume that all the divergence values in the range (*g, i*) are less than *x*, we can conclude that *y*_*h*_[*x* − 1] = 0, ∀ *g < h < i*. This includes *d*_*k*_[*g* + 1] *< x*, which implies that *y*_*g*+1_[*x* − 1] = *y*_*g*_[*x* − 1] = 0. However, we already have the case that *d*_*k*_[*g*] = *x*, which means that *y*_*g*_[*x* − 1] = 1 and *y*_*g−*1_[*x* − 1] = 0. That is a contradiction for the value of *y*_*g*_[*x* − 1] which proves that it cannot be the case that all the divergence values are less than *x*. Hence, there must be a divergence value greater than *x* in the range (*g, i*). □

### 3.2 Finding blocks of starting matches

In a PBWT panel, neighboring haplotypes sharing matching segments cluster together. Such a collection of neighboring matches is called a *block*. Such blocks of matches are separated by a haplotype whose divergence value is greater than the difference of the site being observed and the length threshold specified. When iterating over the divergence array at a certain site, *d*_*k*_, the divergence value properties discussed above restrict the combinations for the ordering of divergence values in the array. This in-turn assists in finding such blocks of matches where the matches share the same starting position.

**Fig. 1** shows an example of using the divergence value properties to find blocks of starting matches of length, *L* ≥ 2. It shows a reverse prefix sorted panel of 10 haplotypes at site *k* = 4, where *d*_4_ shows the values of the divergence array and *a*_4_ shows the prefix array. **Fig. 1(a)** shows the first block of starting matches found and **Fig. 1(b)** shows the second block found. These two blocks are separated by the sequence indexed 3 in *a*_4_ with divergence value 3 which is greater than *k − L* = 2. The grey box at site *k* = 1 in the figures show the two clusters being formed within a block, called *groups*. Within each block, the top group consists of zeros at site *k* − *L* − 1 and the bottom group consists of ones at that site. It’s also to be noted that a block of starting matches must contain only one sequence with divergence value equal to *k − L* value. In the blue block, that sequence is 1, and in the red block, that sequence is 9. The haplotypes belonging to the same block but different groups constitute the actual matches that are of minimum length 2 and start at site *k* = 2. These starting matches are given by the tuples (3, 1), (3, 4) for the blue block and (0, 9) for the red block.

**Figure 1:**
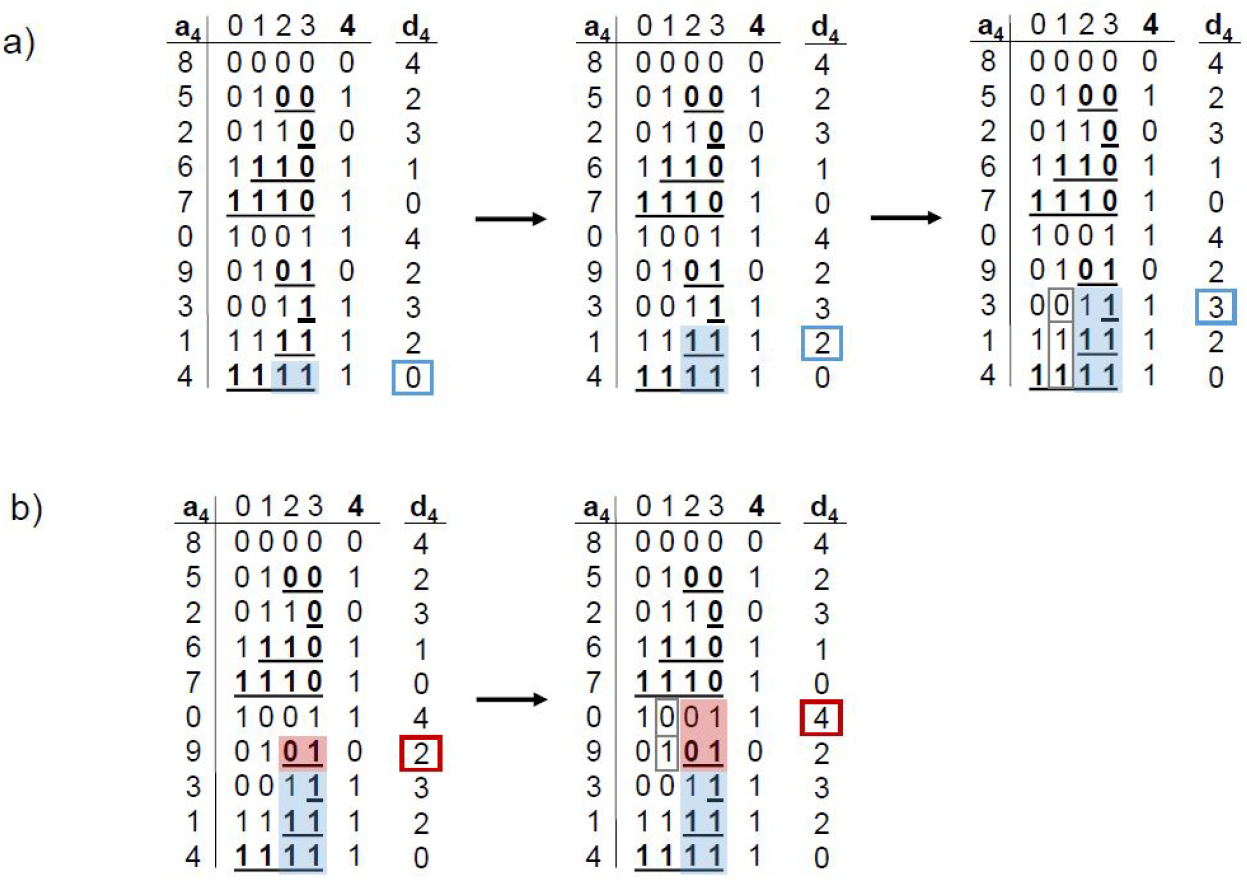
Two blocks of matches of minimum length 2 starting at site *k* = 2 in a panel of 10 haplotypes reverse prefix sorted at *k* = 4. The process of block detection while scanning the divergence array is shown. The rightmost colored rectangle (under *d*_4_) shows the divergence value being scanned while the solid colored rectangles, blue in (*a*) and red in (*b*), show the actual matching blocks. The grey boxes at position *k* = 1 show the two groups within each block.

### 3.3 double-PBWT

*double-PBWT*, as the name suggests, uses two columns of PBWT simultaneously scanning the panel from left to right. The two columns are 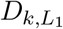 and 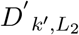 where column 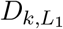 is operating at site *k* finding matches of length at least *L*_1_ ending at site *k* while column 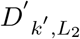 is operating at site *k*′, *L*_2_ sites ahead of *k* such that *k*′ = *k* + *L*_2_. This leading column finds matches of length at least *L*_2_ starting from site *k*. Such a formulation of mcPBWT enables us to evaluate matches flanking on either side of a site. Such flanking matches can be defined by a composite haploytpe match pattern with *B* = 2, namely, double haplotype match pattern where 𝒫 = {*p*_0_, *p*_1_}, |*c*(𝒫)| = 1, *W* (*p*_0_) = *W* (*p*_1_) = 2 and *g* = 0(for simplicity). This form of composite pattern is structured to signify patterns of recombination. Such combination of matching pair segments were also shown to be potential phasing errors[7]. Here, we introduce and focus on one such variation of this composite pattern termed alternating match but the algorithm can find all the other possible variations as well. We define alternating match and the problem statement formally in the following paragraphs.

### Alternating Match Definition

An alternating match is a strict case of a double haplotype match pattern, 𝒫 = {*p*_0_, *p*_1_}, |*c*(𝒫) | = 1, *W* (*p*_0_) = *W* (*p*_1_) = 2 where individual information is also encoded. If we assume the single thread haplotype belongs to an individual say, *A*, the non-thread haplotypes in *p*_0_ and *p*_1_ must be the complementary haplotypes of the same individual, say, *B*. Such a case of double haplotype match pattern is termed as an *alternating match*.For instance, if *p*_0_ is a match between *A*_*i*_[*k*_1_, *k*_2_) and *B*_*j*_[*k*_1_, *k*_2_)where, *i, j* ∈ {0, 1} and *A*_*i*_, *B*_*j*_ are haplotypes of individuals *A* and *B* respectively and *A*_*i*_ is the thread haplotype. Then *p*_1_ must be a match between *A*_*i*_[*k*_3_, *k*_4_) and *B*_*j*′_[*k*_3_, *k*_4_), where *B*_*j*′_ is the complement haplotype of individual *B*. Here, *B* is said to have an alternating match with respect to *A*. **Fig. 2** shows the double haplotype match pattern and an alternating match. **Fig 2(a)** shows the double haplotype match pattern where *H*1, *H*2 and *H*3 are three haplotypes. Here, sequence id set of *p*_0_, i.e., *c*(*p*_0_) = {*H*1, *H*2} and sequence id set of *p*_1_, *c*(*p*_1_) = {*H*1, *H*3} where *c*(*p*_0_) ⋂ *c*(*p*_1_) = {*H*1} is the thread haplotype. This form of double haplotype match represents crossover breakpoints. Similarly, **Fig. 2(b)** shows an example of an alternating match where individual *B* has an alternating match with respect to individual *A*. Here, sequence id set of *p*_0_, i.e., *c*(*p*_0_) = {*A*_0_, *B*_0_} and sequence id set of *p*_1_, *c*(*p*_1_) = {*A*_0_, *B*_1_} such that *A*_0_ is the thread haplotype and *B*_0_ and *B*_1_ are complementary haplotypes of individual *B*. The first segment of the alternating match (the blue segment) is of length *L*_1_ and the second yellow segment is of length *L*_2_. **Fig. 2(b)** shows the case where the boundaries of the two pairs are juxtaposed such that *k*_2_ = *k*_3_, i.e. *g* = 0.

**Figure 2:**
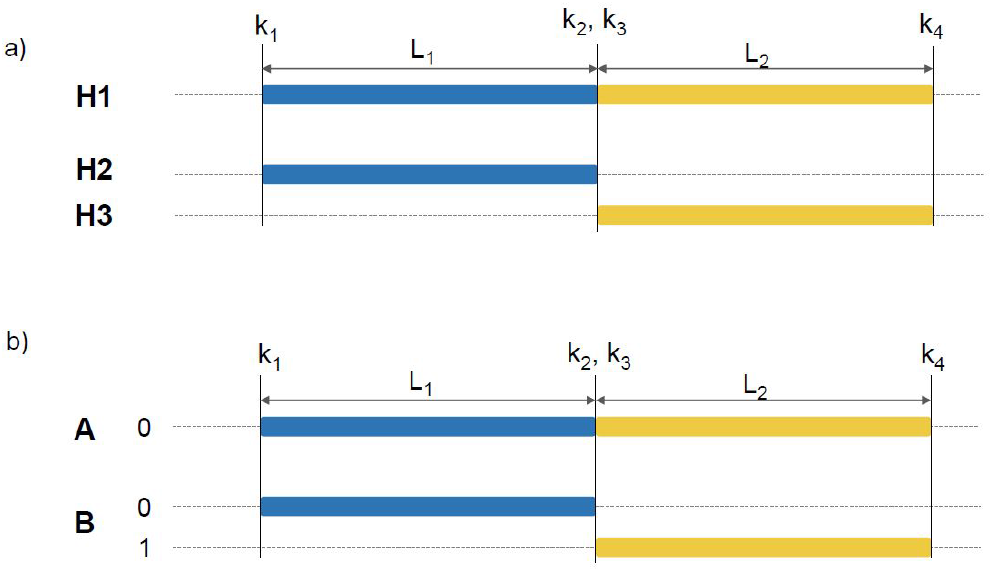
Double haplotype match patterns (a) Double haplotype match pattern where *c*(*p*_0_) = {*H*1, *H*2}, *c*(*p*_1_) = {*H*1, *H*3} and *L*(*p*_0_) ≥ *L*_1_, *L*(*p*_1_) ≥ *L*_2_. Such formation represents crossover breakpoints. (b) Alternating match where *A* and *B* are two individuals and individual *B* has an alternating match with respect to individual *A*.

### Problem Statement

Given a phased haplotype panel and user specified length parameters *L*_1_, *L*_2_ and *g* (here assumed 0 for simplicity), find all the alternating matches. We discuss the simple case of the alternating matches with strict boundaries as shown in **Fig. 2(b)** such that *g* = 0. The algorithm proceeds where the leading PBWT column *D*′ first stores the information on starting matches using block and group properties, passes this information to the lagging PBWT column *D* which then checks if an alternating match exists for each ending match pair found of minimum length *L*_1_. These two PBWT columns are *L*_2_ (*k*′ − *k* = *L*_2_) distance apart so that the starting matches found are of length at least *L*_2_.

**Algorithm 1** shows the working mechanism for PBWT column *D*′. It updates the block and group information for all the starting matches found at a given site. The *block* array of size *M* keeps track of the block-membership of all the haplotypes, where the index of the array represents the haplotypes’ indexes. An integer value (*id*) is assigned to haplotypes belonging to the same block. *group*, an array of size *M* distinguishes between the haplotypes of the block with 0 or 1 at *k*′ *L*_2_ 1 position. The index of this array also represents the haplotypes. *rblock* stores the same block and group information in a dictionary format where it is indexed by the block id to find the double haplotype match pattern show in **Fig. 2(a)**. The divergence values and prefix arrays are calculated using Durbin’s algorithm 1 and 2 [1]. The *block* and *group* arrays along with *rblock* are updated simultaneously as the divergence and prefix arrays are updated so the time complexity to find the blocks of starting matches is *O*(*M*) at each site and *O*(*MN*) for the all sites. The block and group arrays are passed on to the PBWT column *D* where it decides if an alternating match is found. Here, the divergence array is scanned from *M* − 1 to 1 as it’s more intuitive to understand the formation of block and groups but it can be scanned in the other direction without affecting the algorithm as shown later in Algorithm 4

**Algorithm 2** is responsible for finding the other set of matches ending at site *k* and deciding if an alternating match exists. This algorithm scans the panel from site 0 to *N* simultaneously as *D*′ scans the panel ahead of it. It finds all matches ending at site *k* using Durbin’s algorithm 3 [1]. For every such ending match, it checks to see if there’s a starting match that satisfies as an alternating match. This is done using the block and group arrays passed from *D*′. When the condition is met, the alternating match is reported. This checking is done in constant time since array access is constant time. Because of this constant time lookup for every pair of ending match found, the time complexity depends on the number of such match pairs found. We define *C* as the total number of match pairs found across all the sites. Therefore, the time complexity for this column is *O*(*C*).

#### Algorithm 1 Leading PBWT at column 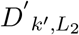: Find blocks and groups of matches starting at position *k*

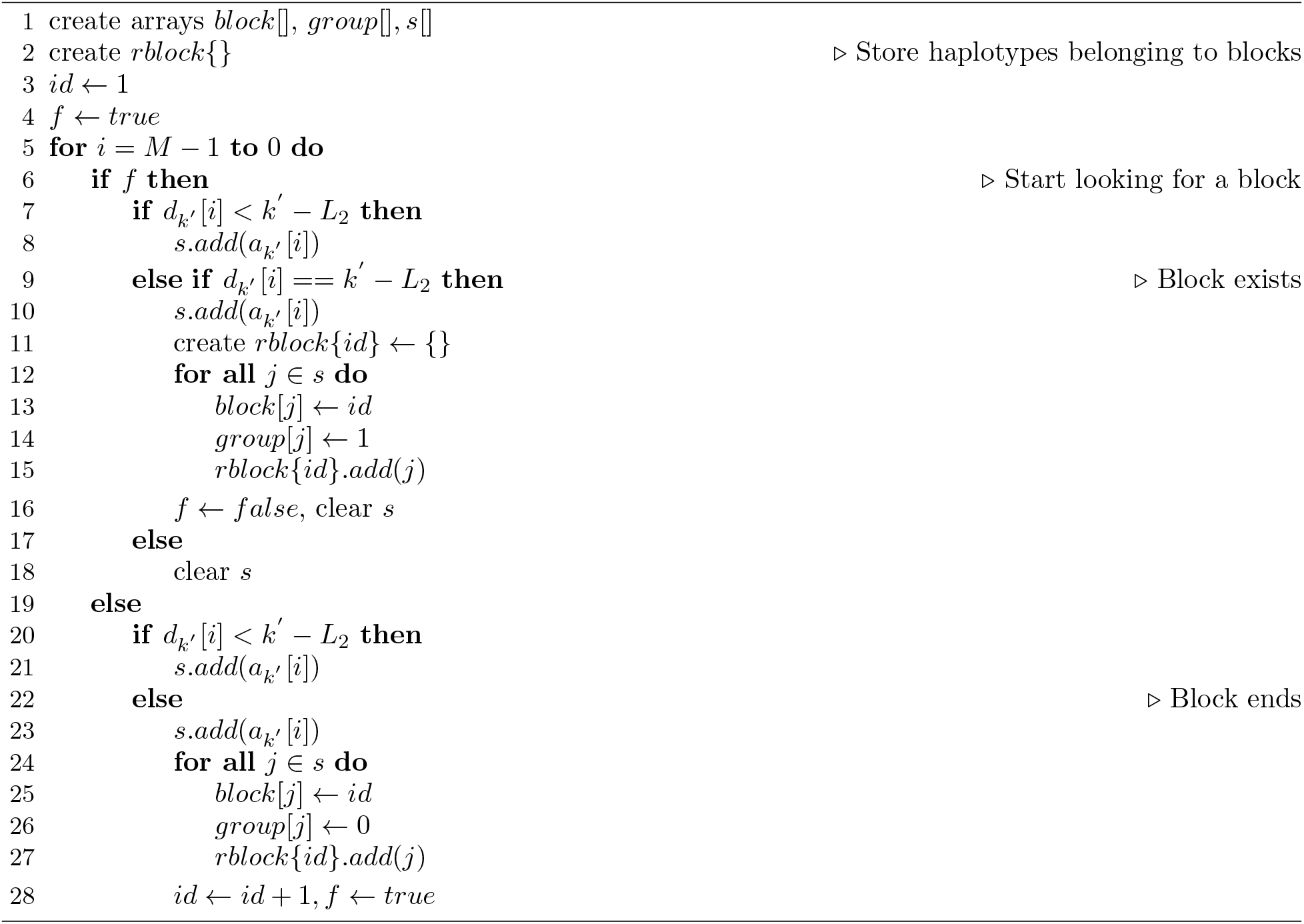

It is to be noted that this algorithm does not handle for the double haplotype match pattern in **Fig. 2(b)** but *rblock* can be used to query such patterns easily. For every ending match pair, like *H*1 and *H*2 detected by lagging PBWT at column *D*, the haplotypes belonging in the blocks of *H*1 and *H*2 are scanned using *block* and *rblock*, to find the second matching segment *H*1 and *H*3 (*H*1 being the thread haplotype) or *H*2 and *H*3 (*H*2 being the thread haplotype). This scanning process is done in *O*(*b*) time for every ending match pair where *b* is the average number of haplotypes in a block. Hence, the overall time compleixty for such an algorithm would be *O*(*C * b*) across all the sites.

**Algorithm 3** shows the synchronous execution of both PBWT columns as it does a one pass scan on the panel. Since both columns move simultaneously across all the sites, the overall complexity of the algorithm is *O*(*MN* + *C*). While we only show the case of alternating match when *g* = 0, these algorithms can be extended to handle for overlaps or gaps (*g* ≥1) by altering the distance between the two PBWT columns.

#### Algorithm 2 Lagging PBWT at column 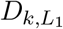 : Report matches of length at least *L*_1_ forming alternating matches

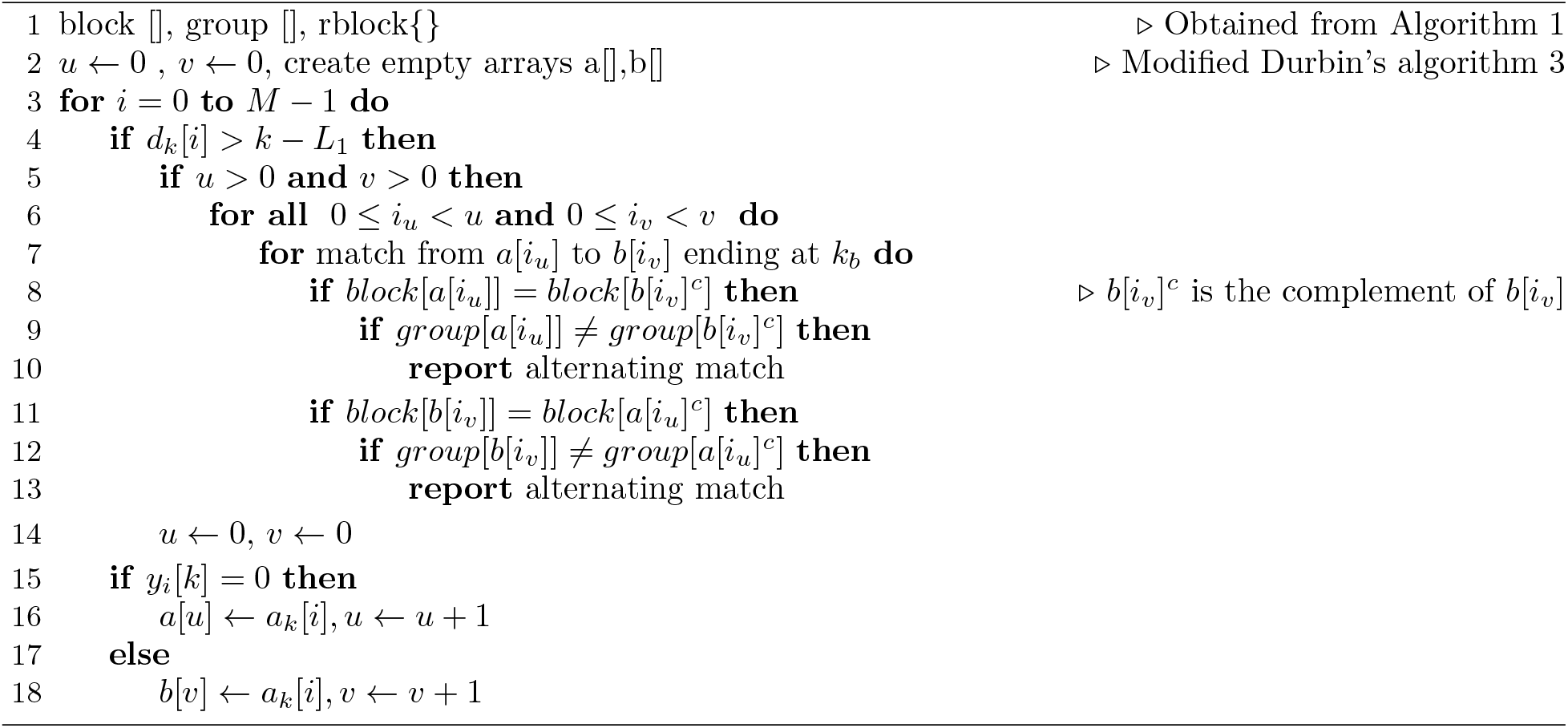

#### Algorithm 3 *double-PBWT*: Simultaneous run of two PBWT columns

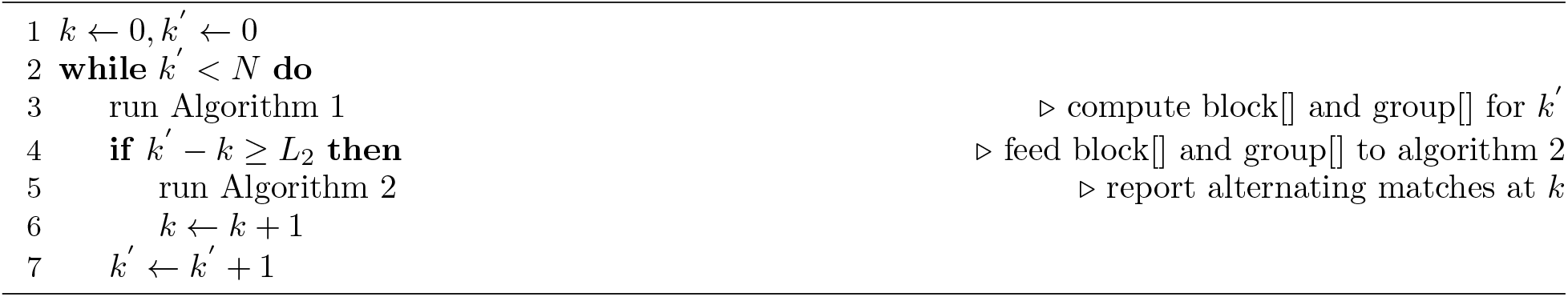

### 3.4 double-PBWT: Comparing Block of Matches

So far we’ve used *double-PBWT* to find double haplotype composite matching patterns where *W* (*p*_*b*_) = 2, *b* = 0, 1 but here we take advantage of its versatility to make comparisons between blocks of matches, i.e. *W* (*p*_*b*_) ≥ 2, *b* = 0, 1. The main idea here is to only evaluate matching blocks found by the two PBWT columns when they satisfy user-specified constraints of a valid block structure. A block structure is defined as a block consisting of at least *W*′ haplotypes in common and sharing at least *L* long segments. This definition is adapted from cPBWT [13] and allows us to process composite match patterns in blocks. Such block-based comparison can be useful in studying recombination patterns too [16]. Here, both columns of PBWT store haplotypes that belong to different blocks and the blocks found by the two columns are compared to see if they share at least *W*′ haplotypes. When this requirement is met, the blocks are output. While **Fig. 3** shows the comparison of blocks of haplotypes, this can also be extended to find alternating matches. Since alternating matches have more structure in terms of individuals that the haplotypes belong to, the algorithm needs to be modified to account for this constraint. For alternating matches in a block structure, additional constraints can be specified for the minimum number of thread haplotypes i.e. |*c*(𝒫)| ≥ *w*_*min*_ and the minimum number of alternating individuals that should be present in a block-structure.

**Fig. 3** shows an example of analyzing blocks of matches of length *L*_1_, *L*_2_ ≥ 2 on either side of site 3. Here, a valid block structure should have at least 2 haplotypes (*W*′ ≥2). The only valid block structure found is shown in the middle with two haplotypes (6, 1) common to the two top blocks indicating that they share an extending match. It can be seen that this can be generalized to handle mismatches to study recombination patterns by adjusting the distance between the two PBWT columns.

**Figure 3:**
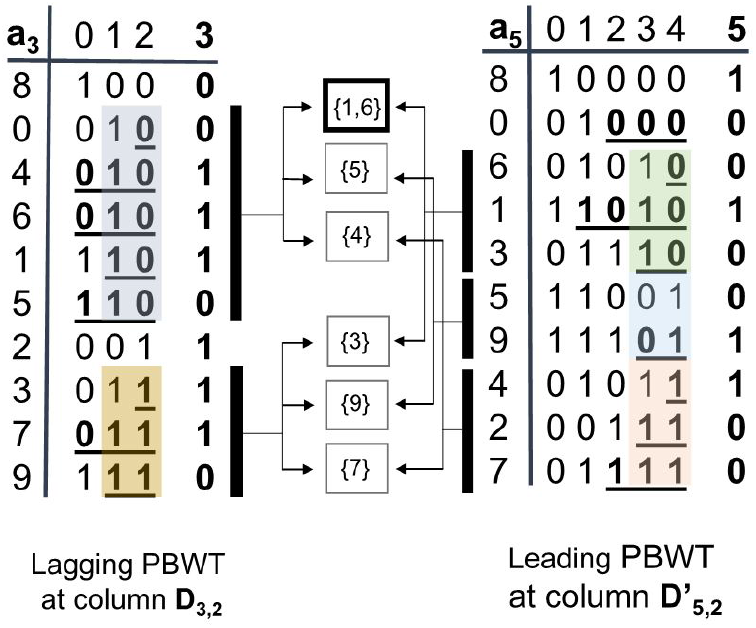
Double PBWT finding blocks of matches of length *L*_1_, *L*_2_ ≥ 2 and *W*′ ≥2. (Left) Haplotype panel with 10 haplotypes reverse prefix sorted at *k* = 3. The two colored boxes represent the block of matches ending at *k* = 3. (Right) Haplotype panel reverse prefix sorted at *k*′ = 5 where the colored boxes represent blocks of matches starting at *k* = 3. (Middle) The boxes in the middle show haplotypes common between the lagging and leading blocks. The grey block from lagging PBWT column *D*_3,2_ and green block from leading PBWT column *D*′_5,2_ share two haplotypes {1, 6} to satisfy *W*′ *≥* 2 showing that the two haplotypes have an extending match.

### 3.5 triple-PBWT

*triple-PBWT* is the case of mcPBWT where three columns of PBWT are utilized. Each column has the freedom to find matches ending at those sites or starting few sites before. Here, we define *triple-PBWT* with three columns 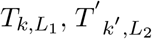 and 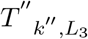 where, *k*′ = *k* +*L*_2_ and *k*^*′′*^ = *k* +*L*_2_+*L*_3_. *triple-PBWT* can be useful in finding a triple haplotype composite match pattern (*B* = 3), 𝒫 = {*p*_*b*_}, |*c*(P)| = 1 and *W* (*p*_*b*_) = 2, *b* = 0 2. The set of length constraints for the three matching pairs are {*L*_1_, *L*_2_, *L*_3_} such that *L*(*p*_0_) ≥ *L*_1_, *L*(*p*_1_) = *L*_2_ and *L*(*p*_2_) ≥ *L*_3_ and *L*_2_ *<< L*_1_, *L*_3_. Here, *L*_2_ is restricted to be a short match in comparison to *L*_1_ and *L*_3_ to emulate a gene-conversion tract. **Fig. 4** shows an example of triple haplotype match pattern. Here, *H*1, *H*2 and *H*3 are three haplotypes where, *H*1 is the thread haplotype. Similarly, the sequence id sets are *c*(*p*_0_) = {*H*1, *H*3}, *c*(*p*_1_) = {*H*1, *H*2} and *c*(*p*_2_) = {*H*1, *H*3}. The three columns of *triple*-PBWT are represented by the vertical dashed lines. Here, column *T* runs a standard PBWT finding matches of length at least *L*_1_ ending at site *k*. The *T*′ column finds matches of exact length *L*_2_ starting at site *k* and ending at site *k* + *L*_2_ and column *T*′′ finds matches of length at least *L*_3_ starting from site *k* + *L*_2_. While a simple composite pattern like the one shown can be representative of gene-conversion tract, it’s not sufficient condition and care has to be taken since the smaller match pair *p*_1_ could end up providing lots of false positives. Additional information has to be incorporated to this formulation to distinguish false positives from true gene-conversion tracts but this shows one potential use case for the algorithm. The first column behaves similar to **Algorithm 1** and can be easily extended from there. The last column runs **Algorithm 2** as in *double-PBWT*. The only change is for PBWT column *T*′. **Algorithm 4** shows how the exact matches can be catalogued for column *T*′. Exact matches satisfy both restrictions of starting and ending matches and hence the algorithm uses ideas of both starting and ending matches to find them. Here, *block* and *group* arrays along with *rblock* serve the same function as in *double-PBWT. rblock* is a dictionary indexed by the block ids that store haplotype indices belonging to such blocks. This dictionary is used to find exact match pairs like *p*_1_. A new array *end* is introduced which keeps track of whether the haplotypes have 0 or 1 in the next variant site. This *end* array is utilized to filter the exact matches from the starting matches. Since, the *end* array is updated along with divergence and prefix arrays, the time complexity for this algorithm at a given site is *O*(*M*). The overall synchronization of the three columns is similar to *double-PBWT* in that columns *T*′ and *T*′′ catalogue the exact matches and starting matches respectively and pass this information in the form of *block, group, rblock* and *end* (for column *T*′) to column *T*. Then, for every ending match pair detected by column *T*, it takes constant time to look up match pairs *p*_2_ but the haplotypes that belong to *H*1’s block have to be scanned for match pair *p*_1_. When those matching pair segments exist, the triple haplotype match is reported.

#### Algorithm 4 *triple-PBWT* at column *T*′ : Find matches that are of exact length *L*_2_ at site *k*′

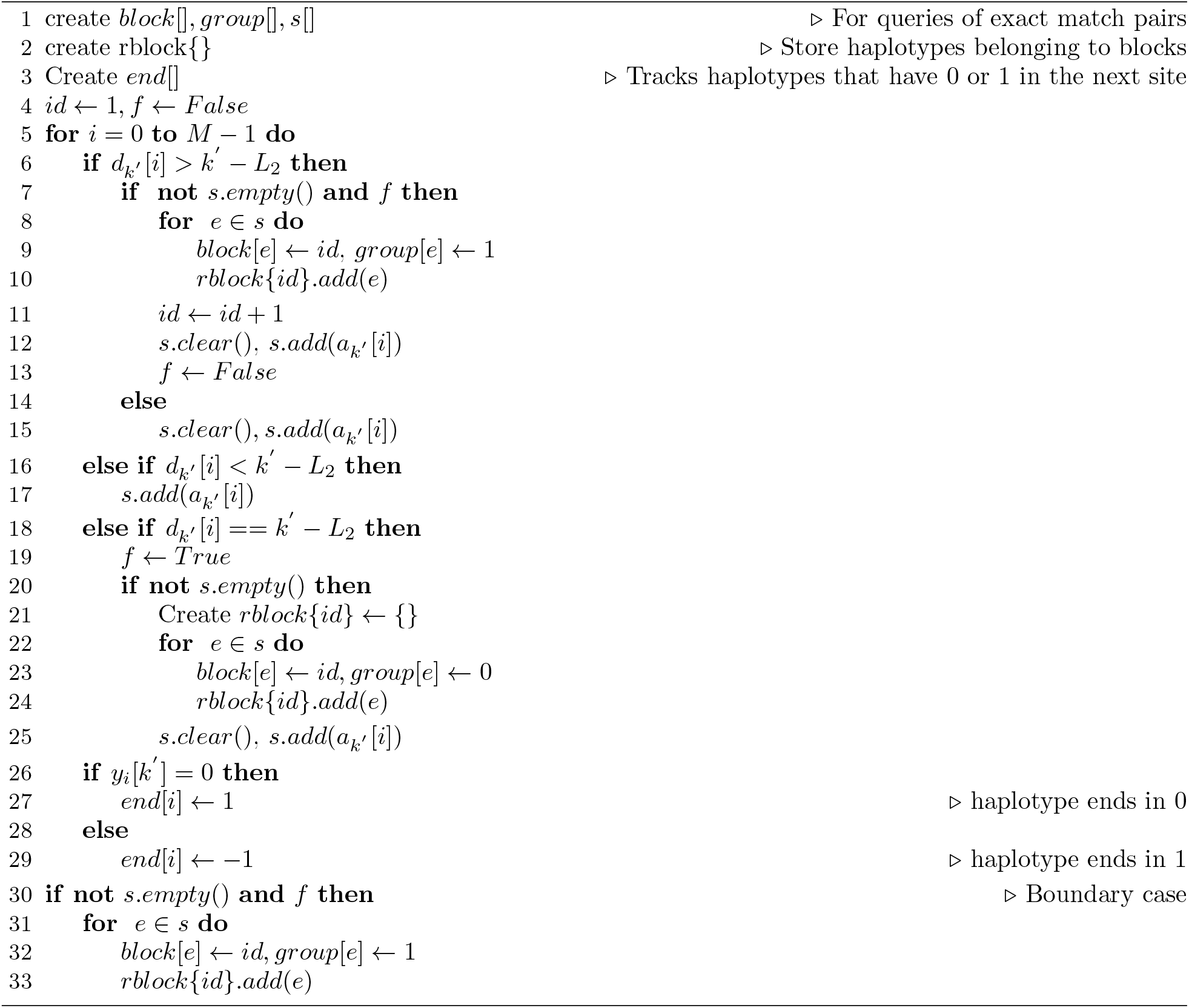

**Figure 4:**
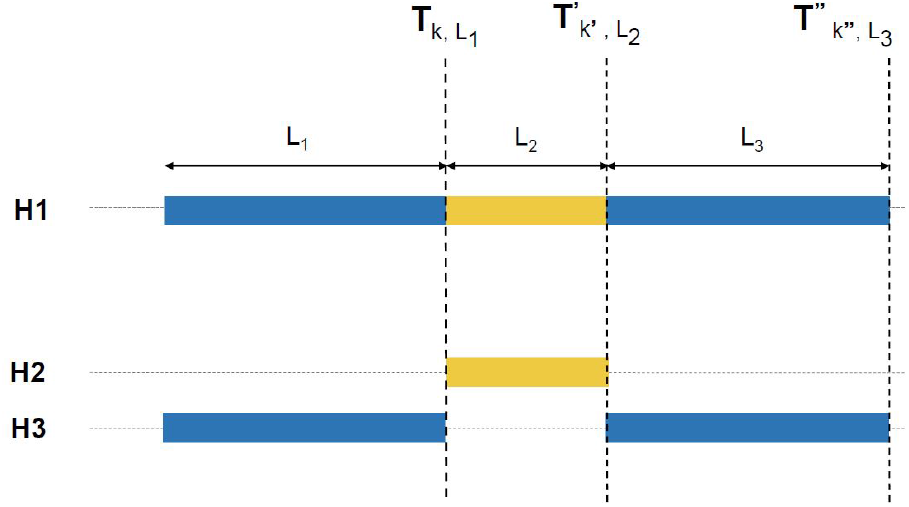
A composite pattern of three matching segments between three haplotypes *H*1, *H*2 and *H*3. This pattern is representative of gene conversion, where the yellow segment is a possible gene conversion tract such that *L*_2_ *<< L*_1_, *L*_3_. The vertical dashed lines show the three simultaneous PBWT runs each operating at sites *k, k*′ and *k*^*′′*^, where *k*′ = *k* + *L*_2_ and *k*^*′′*^ = *k* + *L*_2_ + *L*_3_.

Of the three PBWT columns of *triple-PBWT*, column *T*′′ is the same as the leading column of *double-PBWT* and hence has a time complexity of *O*(*M*) at each site and *O*(*MN*) across all variant sites. For the middle column *T ′*, the time complexity to catalogue the exact matches is *O*(*M*) at each site and *O*(*MN*) across all the sites. It’s important to note that querying of exact matches of length *L*_2_ can be done by accessing *block, group, end* arrays in constant time. Lastly, column *T* makes constant time query for every ending match found to see if the last match pair exists but has a time complexity of *O*(*b*) to find the middle match pair, where *b* is average number of haplotypes in a block. Since, *T* is our trigger column, the time complexity depends on the number of match pairs found by PBWT column *T*. Hence, we define the complexity of this column across all sites similar to *double-PBWT* as *O*(*C* ∗ *b*), where *C* is the total number of match pairs found across all the variant sites. The overall time complexity of Triple PBWT is then *O*(*C* ∗ *b* + *MN* + *MN*), i.e. *O*(*MN* + *C* ∗ *b*).

## 4 Discussion

In this work, we present a more flexible and powerful variation of PBWT for detecting composite haplotype matches. The original formulation in PBWT for the haplotype matching problem only captures the matching pattern at every single column separately. Our algorithm, however, simultaneously captures the patterns across multiple columns of PBWT. In the single column matching formulation, at the active column of the PBWT, one only has access to the information in the past but is uninformed about future columns. In our formulation, the columns at the forefront can provide “look ahead” information allowing the algorithm to make complex decisions. Our flexible algorithm’s capability to analyze composite matching patterns opens new potentials of the PBWT data structure.

The proposed method does not require to output or book keep the matches which will be very useful in analyzing large haplotype panels with millions of individuals. While the PBWT algorithm is able to find all matches efficiently, the number of matches in large cohorts may be enormous. As a result, analyzing composite patterns like alternating matches after the PBWT run may not be very efficient. Hence, a flexible algorithm like *double-PBWT* would be useful in such cases. Additionally, we also showed that the *triple-PBWT* could find composite haplotype match representative of gene-conversion tracts. This shows that mcPBWT has potential to allow and adjust for flexible matching criteria and is suitable for more general-purpose settings.The double haplotype match pattern discussed where *H*1 is the thread haplotype and *H*2 and *H*3 have matches with it not only identifies more recombination events but also provides plausible evidence that *H*2 and *H*3 coalesce more recently. This could help to determine the time of the recombination events, and also help “triangulating” the genealogical relationship among individuals carrying these matching segments. Such analyses can also be conducted using blocks to enable stronger signal using mcPBWT.

## Acknowledgements

PS, AN, DZ and SZ were supported by the National Institutes of Health grant R01 HG010086. AN, DZ and SZ were also supported by the National Institutes of Health grants R56 HG011509. AN and DZ were also supported by the National Institutes of Health grant OT2-OD002751.

## Notes

### Competing Interest Statement

The authors have declared no competing interest.

